# Reconstructing human early embryogenesis in vitro with pluripotent stem cells

**DOI:** 10.1101/2021.03.12.435175

**Authors:** Berna Sozen, Victoria Jorgensen, Meng Zhu, Tongtong Cui, Magdalena Zernicka-Goetz

**Affiliations:** Plasticity and Self-Organization Group, Division of Biology and Biological Engineering, Caltech, Pasadena, CA 91125; Mammalian Development and Stem Cell Group, Department of Physiology, Development and Neuroscience, University of Cambridge, Cambridge CB2 3EG, UK; Department of Genetics, Yale School of Medicine, Yale University, New Haven, CT 06520; Genetics Department, Harvard Medical School, Harvard, Boston, MA 02215

## Abstract

Understanding human development is of fundamental biological and clinical importance. Yet despite its significance, insights into early developmental events in humans still remain largely unknown. While recent advances show that stem cells can mimic embryogenesis^1–9^ to unravel hidden developmental mechanisms, a stem cell-based model of early human embryogenesis is lacking. Here, we use human extended pluripotent stem cells^10^to reconstitute early human development in 3-dimensions and recapitulate early embryo-like events. We first perform a systematic characterisation to reveal unique signalling requirements for building the human pre-implantation blastocyst. Further, we show that these *in vitro* stem cell-derived blastocyst-like structures are able to undertake spatiotemporal self-organisation to mimic peri-implantation remodelling in which a polarised rosette opens up the amniotic cavity within a developing disc. The hallmarks of human early development displayed by this stem cell-based *in vitro* model mimics features of embryonic day 3 to day 9/10 of natural development. Thus, this platform represents a tractable model system to contribute to the basic understanding of cellular and molecular mechanisms governing early embryonic events in humans and to provide valuable insights into the design of differentiation protocols for human stem cells in clinical applications.

---

Development begins in the zygote, a single fertilised egg. While the undifferentiated zygote holds the potential to form all embryonic and extra-embryonic lineages^11,12^, cells gradually lose potency as they diversify into specialized functional cell types as development progresses. During the acquisition of cell fate, cells follow a precise sequence of regulative events, governed by distinct mechanisms that maintain a fine balance between pluripotency and differentiation. The first cell fate decisions take place after embryonic day 3 (D3) in the human pre-implantation embryo and generate spatially segregated lineages by D5/6, forming an organised, hollow structure referred to as the blastocyst **(Fig. 1a)**. The blastocyst initially comprises two distinct cell populations: the extra-embryonic trophectoderm (TE), an outer epithelium, and the embryonic pluripotent inner cell mass (ICM), positioned to one side of the cavity **(Fig. 1a)**. The ICM is considered to represent an initial multi-lineage priming state, which undergoes a second differentiation event just prior to implantation at D6-7^13^. During this event, the transient ICM gives rise to both the epiblast (Epi), the source of all foetal cell lineages, and the extra-embryonic hypoblast (Hypo) **(Fig. 1a)**. Following implantation, the ICM-derived lineages undergo a series of morphological changes leading to the formation of a disc-shape structure.

**Figure 1.**
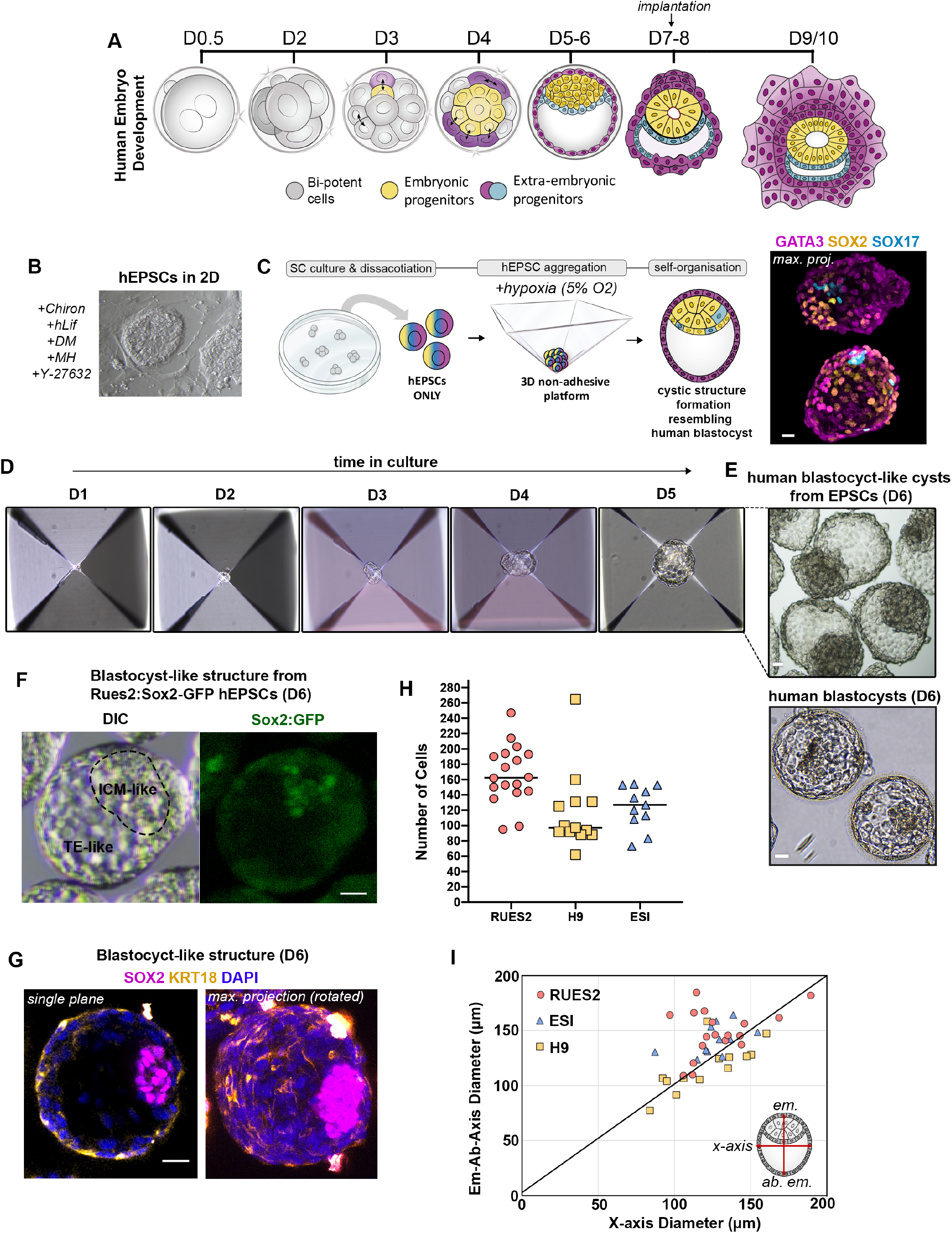
A 3D system to mimic early human embryogenesis from hEPSCs. **A.** Scheme for natural human pre/peri-implantation embryo development. **B.** A representative dome-shaped naïve pluripotent hEPSC colony in 2D culture. **C. Left:** Schematic of an AggreWell and aggregation protocol to mimic early human development solely with hEPSCs. **Right:** Representative examples of cystic structures generated from a typical experiment after 4 days demonstrate three lineages, resembling blastocyst stage natural embryo. Lineage markers: SOX2, yellow (Epi-like); GATA3, magenta (TE-like), and SOX17, cyan (Hypo-like). n=10 experiments. **D.** Representative phase-contrast images of hEPSC multicellular aggregates in AggreWell at the indicated time points during 3D culture. **E.** Phase-contrast images of human blastocyst-like structures at D6. (top) and human blastocysts at D6 (bottom) **F.** A representative cystic structure generated from RUES2:SOX2-GFP reporter hEPSC line. **G.** A representative structure immunostained for SOX2 in magenta, KRT18 in yellow to label Epi-like inner compartment and TE-like outside epithelium, respectively. DAPI is shown in blue. Maximum projection image is shown on the right. n=50 structures, 3 experiments. **H.** Quantification for cell numbers in individual cystic structures generated from three established hEPSC lines, ESI0017 (n=11), RUES2 (n=18) and H9 (n=13). **I.** Measurements of axial diameters in cystic structures from ESI0017 (n=11), RUES2 (n=18) and H9 (n=13). Illustration on right shows the two axes measured. All scale bars in the figure indicate 20 um.

Recent *in vitro* culture methods have advanced our abilities to study aspects of natural human embryo development ex-utero^14–16^. These systems, while valuable, have restricted use in investigations of the basic biology of early human embryogenesis because of technical and ethical limitations that hinder significant progress^17^. Thus, the development of methods to recapitulate the principles of development *in vitro* with stem cells is gaining momentum leading to models that successfully recapitulate various aspects of natural embryogenesis with stem cells in mice^2–4,7,9,18–22^ and human^1,23,24^. While these platforms provide great potential for understanding hidden developmental mechanisms in mammals, an *in vitro* model that recapitulates distinct spatiotemporal events in early human embryogenesis remains lacking.

Recent studies showed that pluripotent stem cells (PSCs) can be reprogrammed to a molecular state similar to that of earlier blastomeres with an enhanced developmental potency for both embryonic and extra-embryonic cell lineages, termed extended or expanded pluripotency (EP)^10,25,26^. The impetus to reconstitute the unique development of the early human embryo with stem cells *in vitro* led us to hypothesize that human pluripotent stem cells (hPSCs) under conditions of EP should be able to undertake early cell fate decisions and self-organise into 3-dimentional (3D) embryo-like structures. After conversion of hPSCs to hEPSCs over 5 passages (see Materials and Methods), we confirmed that such cells could acquire the morphological features characteristic of pluripotent cells in the naïve state of pluripotency, including dome-shaped colony formation, as supported by earlier observations^10^ **(Fig. 1b)**. It is noteworthy that we could observe flat cell colonies, a morphological feature characteristic to pluripotent cells at the primed state, in different ratios after each passage, indicating a mixed population of cells at different pluripotent states under EP culture conditions **(Extended Data Fig. 1a)**. Using the multi-inverted-pyramid microwell-based 3D culture system^2,3^, we seeded hEPSCs in small numbers (5-6 cells per microwell) to enable their aggregation and subsequent self-organisation **(Fig. 1c).** Importantly, we developed a culture media that enabled us to generate cystic structures that capture the multi-potency of hEPSCs to generate embryo-like lineages over 4 days of 3D culture (see Materials and Methods). To do this, we first observed that the *in vitro* media normally used for the culture of natural human pre-implantation embryos promoted the formation of these cystic structures (**Extended Data Fig. 1b,** see Methods). In order to support the maintenance of pluripotency and to promote TE differentiation, we mixed 2 parts of this medium with 1 part of *EP*^10^ and 1 part of *hTSC*^27^, two different stem cell-base media (see Materials and Methods). Moreover, we found that low oxygen tension (5% O2) facilitated efficient formation of cavitated structures **(Extended Data Fig. 1c)**, similar to our previous observations of mouse blastoid formation from mouse EPSCs^3^ and in the development of natural human blastocysts^28^.

We next screened different growth factors, cytokines, and small molecules to identify conditions enabling the generation of structures closely resembling human blastocysts with both cavity and early-lineage specification **(Extended Data Fig. 2a-Ee)**. We found that a combination of BMP4 (20ng/ml), WNT antagonist CHIR99021 (2uM), FGF2 (40ng/ml) and ROCK inhibitor Y-27632 (5uM) during the first 48h of 3D culture enhanced cell survival and promoted formation of compact cellular aggregates **(Extended Data Fig. 2a-f)**. Additionally, we pulsed the cells with ALK5 inhibitor A83-01 (2uM) to promote TE differentiation^27^ for the first 48h of 3D culture. After this initial 48h, A83-01 was omitted from the culture media in order to avoid a complete loss in pluripotency. Concomitantly, the concentration of FGF2 was decreased by half (20ng/ml) for the same purpose^1^. Using this optimized condition, we consistently observed the emergence of cavitated cystic structures, 3 to 4 days after cell seeding **(Fig. 1d).** By day 6 of 3D culture, the structures exhibited typical blastocyst-like morphology, forming a cohesive single outside layer, with an enlarged cavity, and an internal acentric compartment **(Fig. 1d-e)**. Importantly, when SOX2:GFP reporter EPSCs^29^ were used to generate these structures, we found that the ICM-like compartment was positive for GFP signal, while TE-like outer cells were negative **(Fig. 1f)**. We also detected KRT18 expression in the outer layer, confirming correct lineage specification and resemblance to natural blastocysts^13^ **(Fig. 1G)**. The average cell number and diameter of these structures was comparable to those of human blastocysts^30^ **(Fig. 1h, i).** To highlight the robustness of the protocol, all results were repeated with three established hPSC lines, RUES2, ESI-0017 and H9, upon their conversion to EPSCs **(Fig. 1g, h)**.

**Figure 2.**
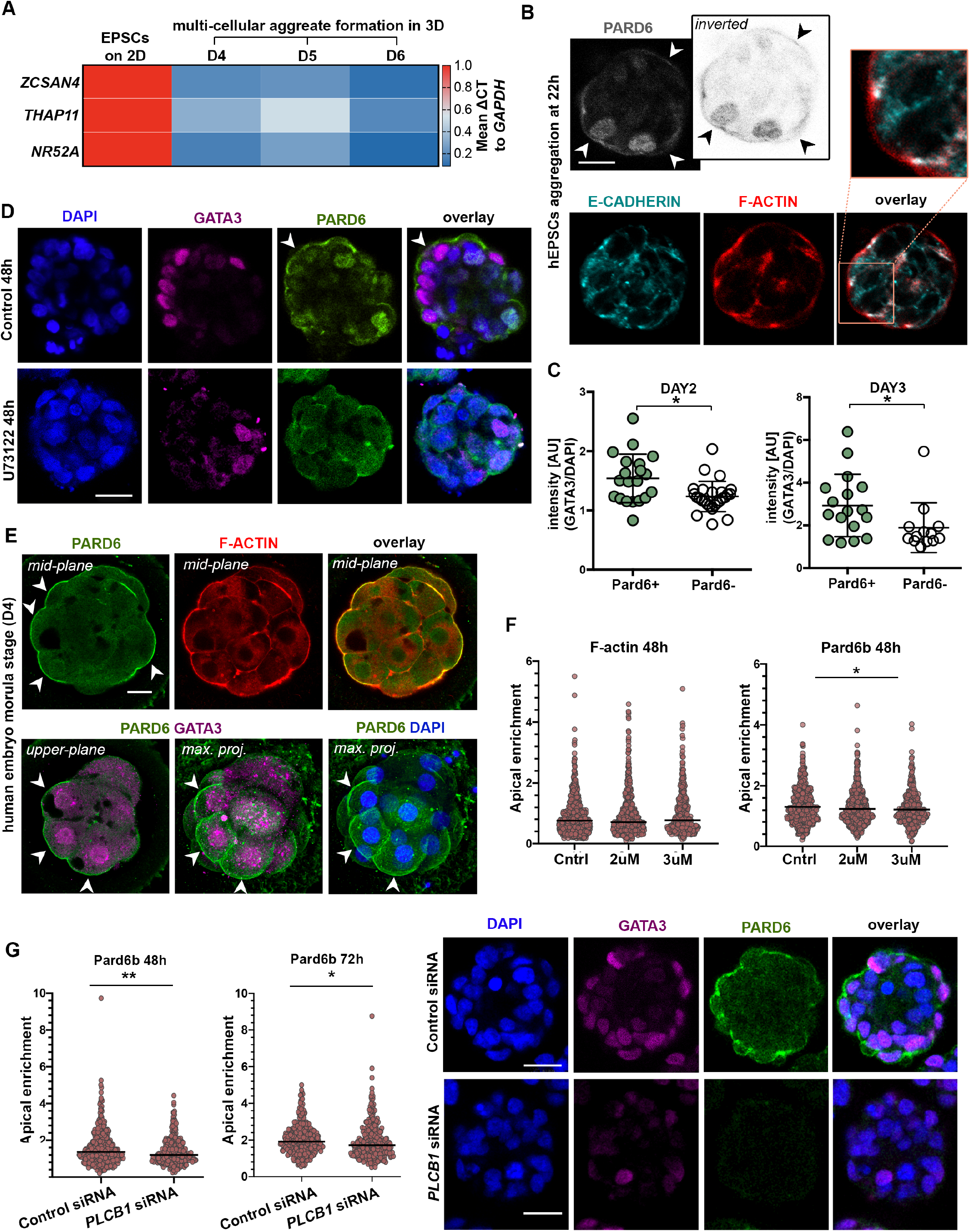
Recapitulation of key pre-Implantation developmental processes. **A.** Bulk qRT-PCR analysis of genes *ZCSAN4, THAP11, NR52A* in EPSCs in 2D, and multicellular aggregates at day 4, 5, 6 formed in 3D represented as a heatmap of global ΔΔCt to *GAPDH.* 20 multicellular cystic structures were pooled per group from 3D culture and a minimum of 10K hEPSCs were collected from the 2D culture. **B.** Immunostaining of hEPSC aggregates at 22h for PARD6 (grey), F-ACTIN (red) and E-CADHERIN (cyan). n=300 aggregates, 3 experiments. **C.** Quantification of GATA3 expression in cells with or without PARD6 apical enrichment observed in cells within Day 2 and Day 3 of multicellular aggregates. All measurements normalized to DAPI. Student’s t-test; p<0.05; 3 experiments. Error bars represent S.E.M. **D.** Immunostaining of control and U73122-treated hEPSC aggregates at 48h for PARD6 (green) and GATA3 (magenta). n=300 aggregates, 3 experiments. **E.** A representative natural human embryo at morula stage (D4) stained for PARD6 (green), F-ACTIN (red), GATA3 (magenta). White arrowheads indicate apical PARD6 enrichment in the polarised cells with nuclear GATA3 expression. DAPI is shown in blue. **F.** Apical enrichment quantification of F-ACTIN and PARD6b at 48h in multicellular structures with or without addition of PLC inhibitor (U73122). Control groups received no inhibitor, while the two experimental groups were treated with either 2uM or 3uM U73122. Each dot represents one analysed cell. *p<0.05, Kruskal-Wallis test with a multiple comparisons test. Data is shown as mean S.E.M. n= 3 experiments. Also see Extended Data Figure 3b. **G. Left:** Quantification of Pard6b apical enrichment at 48h and 72h in structures treated with either control siRNA or *PLCB1* siRNA. Each dot represents one analysed cell. *p<0.05, Student’s t-test. Data is shown as mean S.E.M. n= 3 experiments. **Right:** Immunostaining of GATA3 (magenta) and PARD6 (green) in structures treated with either control siRNA (top) or *PLCB1* siRNA (bottom). DAPI is shown in blue. All scale bars in the figure indicate 20 um.

In order to understand if our system captured human-specific regulatory processes of cellular differentiation in early development, we first analysed hEPSCs before and after multicellular aggregate formation at different stages using quantitative RT-PCR (qRT-PCR) **(Fig. 2a)**. We found that molecular markers known to be highly expressed in 2-to-8-cell blastomeres including *ZSCAN4*, *THAP11* and *NR5A2*^10,26,31^ were highly expressed in hEPSCs before aggregation, indicating the potential capacity of these cells to develop into various cell types **(Fig. 2a)**. As expected, upon multicellular aggregate formation these markers became downregulated, suggesting early embryo-like cellular differentiation **(Fig. 2a).** Previous studies have demonstrated that the first lineage segregation event begins with compaction and cell polarisation in the mouse embryo at the 8-cell stage^32–34^. This process remains unclear in human early embryogenesis and only recently have studies begun to shed light on these events^35,36^. Hence, we utilised our platform to analyse the establishment and dynamics of compaction and cell polarisation at the early time-points of multicellular aggregate formation. The assembly of intercellular junctions is characterised by the formation of baso-lateral expression of E-CADHERIN, suggesting that the compaction events are recapitulated in this *in vitro* system within the first 24h of cell aggregation **(Fig. 2b)**. We also found distinct apical enrichment of F-ACTIN **(Fig. 2b)**, with PARD6 enrichment at the apical surface within the first 48h of cell aggregation **(Fig. 2b)** indicative of cell polarisation. Next, we analysed spatiotemporal expression of the transcription factor GATA3, known as the earliest marker of TE specification in human embryogenesis. GATA3 was localised in the nucleus within both polarised and non-polarised cells at day 2 and day 3 of 3D culture, although the intensity of nuclear GATA3 staining was significantly higher in polarised cells, as judged by apical enrichment of PARD6 **(Fig. 2c, d)**. These results correlated with natural human embryos at the morula stage^36^ **(Fig. 2e).** We have recently shown that the PLC-Protein Kinase C (PKC) pathway controls cellular polarisation at early stages of human development^36^. Of note, when U73122 (a selective PLC inhibitor) was applied at 2 μM and 3 μM in hEPSC 3D cultures, there was significant reduction of the nuclear GATA3 signal intensity **(Fig. 2d, Extended Data Fig. 3a)**. This correlated with a decrease in the apical enrichment of PARD6 upon U73122 treatment in 3D culture **(Fig. 2f, Extended Data Fig. 3b).** Finally, we confirmed the relationship between polarisation and outer cell commitment by using siRNA transfection to knock-down (KD) *PLCB1* in order to deplete PLC activity^36^ in cells during 3D aggregation **(Fig. 2g)**. Again, this revealed a significant reduction in both PARD6 and GATA3 expression in hEPSC aggregates at day 3 **(Fig. 2f-g, Extended Data 3c)**. In accord with recent findings in the human embryo^36^, our results further support the role of the acquisition of apicobasal polarity in promoting the expression and nuclear localisation of GATA3 to drive TE specification in early human development.

**Figure 3.**
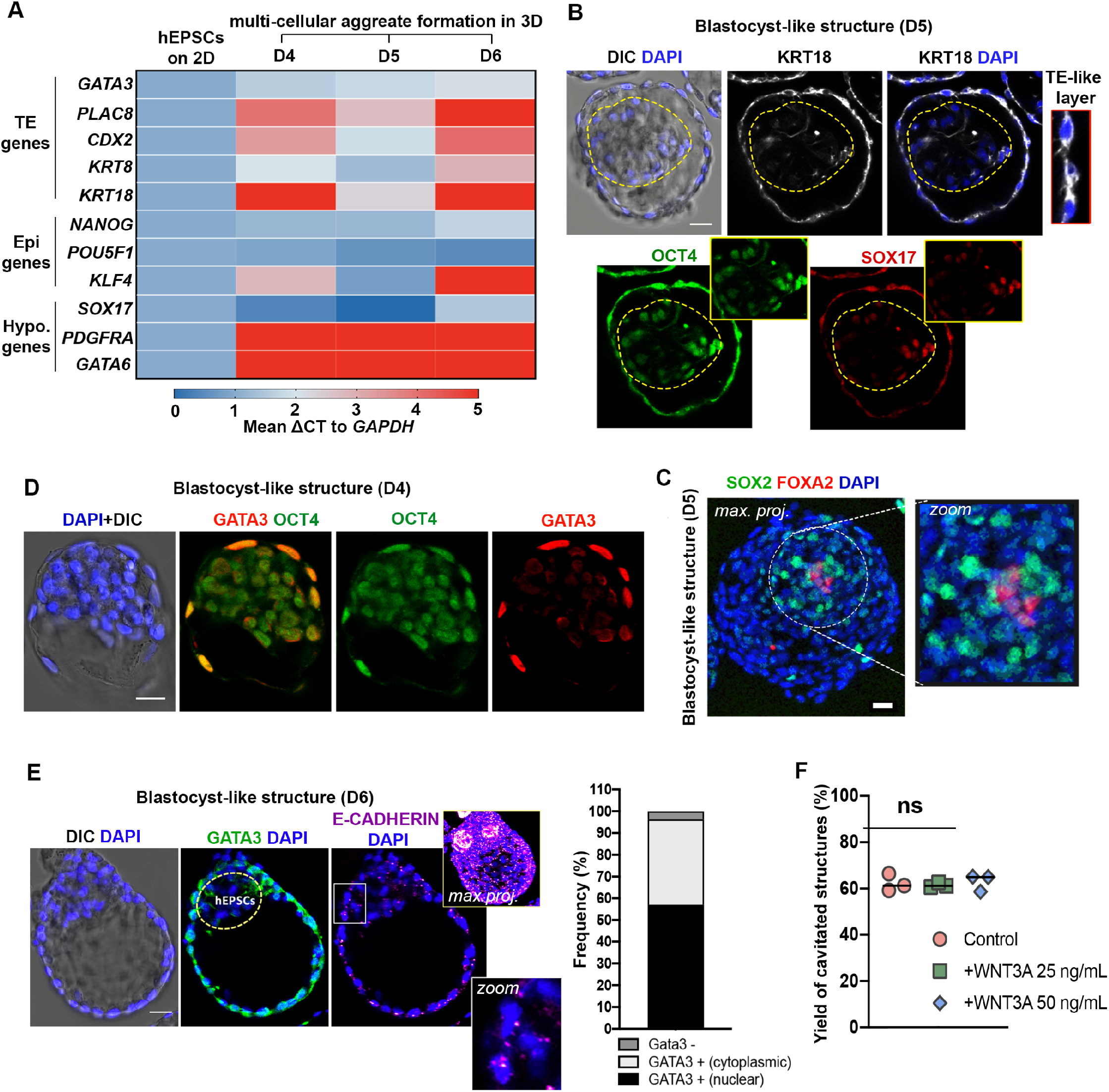
Specification of blastocyst lineages. **A.** Bulk qRT-PCR analysis of blastocyst lineage marker genes in EPSCs in 2D, and multicellular aggregates at day 4, 5, 6 formed in 3D represented as a heatmap of global ΔΔCt to *GAPDH.* 20 cystic structures were pooled per group from 3D culture and a minimum of 10K hEPSCs were collected from the 2D culture. **B.** Immunofluorescence staining of blastocyst-like structures generated from hEPSCs at day 5 for OCT4 (green), KRT18 (white) and SOX17 (red). Zoom image on the right shows TE-like cells with KRT18 expression. DAPI is shown in blue. n= 30 structures, 3 experiments. **C.** A representative blastocyst-like structures generated from hEPSCs at day 5 stained for SOX2 (green) and FOXA2 (red) to reveal Epi/Hypo-like inner compartment (zoom on the right). Image presented as maximum projection. n=20 structures, 2 experiments. **D.** Immunofluorescence staining of blastocyst-like structures generated from hEPSCs at day 4 for OCT4 (green) and GATA3 (red). DAPI is shown in blue. n= 10/23 structures, 2 experiments. **E. Left:** Immunofluorescence staining of GATA3 (green) and E-CADHERIN (magenta) in a representative blastocyst-like structure at day 6. **Right:** Quantification shows frequency of structures at day 6 of 3D culture showing Gata3 nuclear expression (57.03%, 77/135 structures scored); Gata3 cytoplasmic expression (39.25%, 53/135 structures scored); no detectable Gata3 expression (3.70%, 5/135 structures scored). **F.** Quantification shows frequency of cavitated structures in control and WNT3A-supplemented culture. WNT3A is applied in either at 25 or 50 ng/mL concentration. ns, not significant. All scale bars in the figure indicate 20 um.

We next sought to investigate the formation of blastocyst lineages upon cavitation of hEPSC-derived structures from day 4 onwards. We first used qRT-PCR to examine the expression level of core transcription factors responsible for establishing the human blastocyst lineage identity **(Fig. 3a)**. This revealed that genes involved in TE specification, including *PLAC8, CDX2, KRT8* and *KTR18,* were strongly induced upon formation of cystic structures although *GATA3* showed only a marginal increase compared to other molecular determinants of TE identity **(Fig. 3a)**. As expected, crucial transcription factors required for pluripotent Epi specification, including *NANOG* and *POU5F1,* showed similar levels of expression in cystic structures as in hEPSCs cultured in 2D whereas KLF4 was significantly upregulated in the cystic structures **(Fig. 3a).** Finally, we found that the expression of the core Hypo lineage determinant genes, *PDGFRA* and *GATA6*, was highly enriched in cystic structures although *SOX17* did not follow this trend **(Fig. 3a)**. In order to confirm these results spatially and on a protein level, we performed immunofluorescence analysis with well-known lineage markers. In accord with the qRT-PCR results, we observed enrichment for KRT18 in the outside TE-like layer, and expression of OCT4/SOX17 in the Epi/Hypo-like inner compartment **(Fig. 3b).** We further confirmed the Epi/Hypo-like lineage specification with a second set of markers, SOX2/FOXA2^13^ **(Fig. 3c)**. At day 4, some structures displayed constitutive expression of GATA3 in the outer cell layer while maintaining expression of the hPSC/Epi marker OCT4 (10/23 structures scored) **(Fig. 3d).** At later time-points in culture (D6), some structures maintained GATA3 expression in the TE-like outside layer, although this enrichment became mostly cytosolic rather than nuclear (53/135 structures scored) **(Fig. 3e)**. Such late-structures also showed poor expression of E-CADHERIN at day 6 **(Fig. 3e)**. This likely indicates a deficiency in junction assembly during the late cavitation process and may explain the compromised expression of some TE-specific markers as *in vitro* development progresses. WNT3A supplementation is suggested to promote cavitation and thus TE-like lineage identity in *in vitro* mouse blastoid formation^7^, which correlates with canonical WNT expression in the TE lineage during mouse blastocyst development^37^. When we tested this possibility in our human platform, we found that WNT3A addition to the culture media did not make a significant difference in yield of cavitated structures at day 6 **(Fig. 3f)**. This suggested that WNT3A functions differently in human cells/development than in mouse, supporting previous claims^5,38^. Taken together, we conclude that although the transcriptional machinery for blastocyst lineage programming is robustly initiated in hEPSCs, a continuum between distinct cell fates then develops, suggesting a failure in full trans-differentiation.

We next tested the developmental capacity of these hEPSC-derived blastocyst-like structures to develop beyond implantation stages by culturing them in our previously established human embryo *in vitro* culture (IVC) platform^15^ **(Fig. 4a)**. Within 24h in IVC, the blastocyst-like structures reorganised into post-implantation-like disk-shaped structures, 60% of which had a SOX2 positive Epi-like inner compartment surrounded by a KRT18 and GATA3 positive extra-embryonic-like compartment **(Fig. 4b-d)** with some structures also specifying FOXA2-expressing Hypo-like cells **(Fig. 4c)**. Significantly, we found that within 24h in IVC, SOX2 positive cells in the Epi-like inner compartment became radially organised around a small central lumen **(Fig. 4d)**. The formation of a small lumen was confirmed by PODXL expression **(Fig. 4e)**. This demonstrates that this hEPSC-based system can undertake one of the hallmarks of human Epi lineage transformation at implantation, namely the acquisition of a polarised epithelium and the formation of a lumen, the prospective amniotic cavity^14–16^. This strongly suggests that this hEPSC-derived system is undertaking the cell rearrangements that characterize the initiation of post-implantation human morphogenesis.

**Figure 4.**
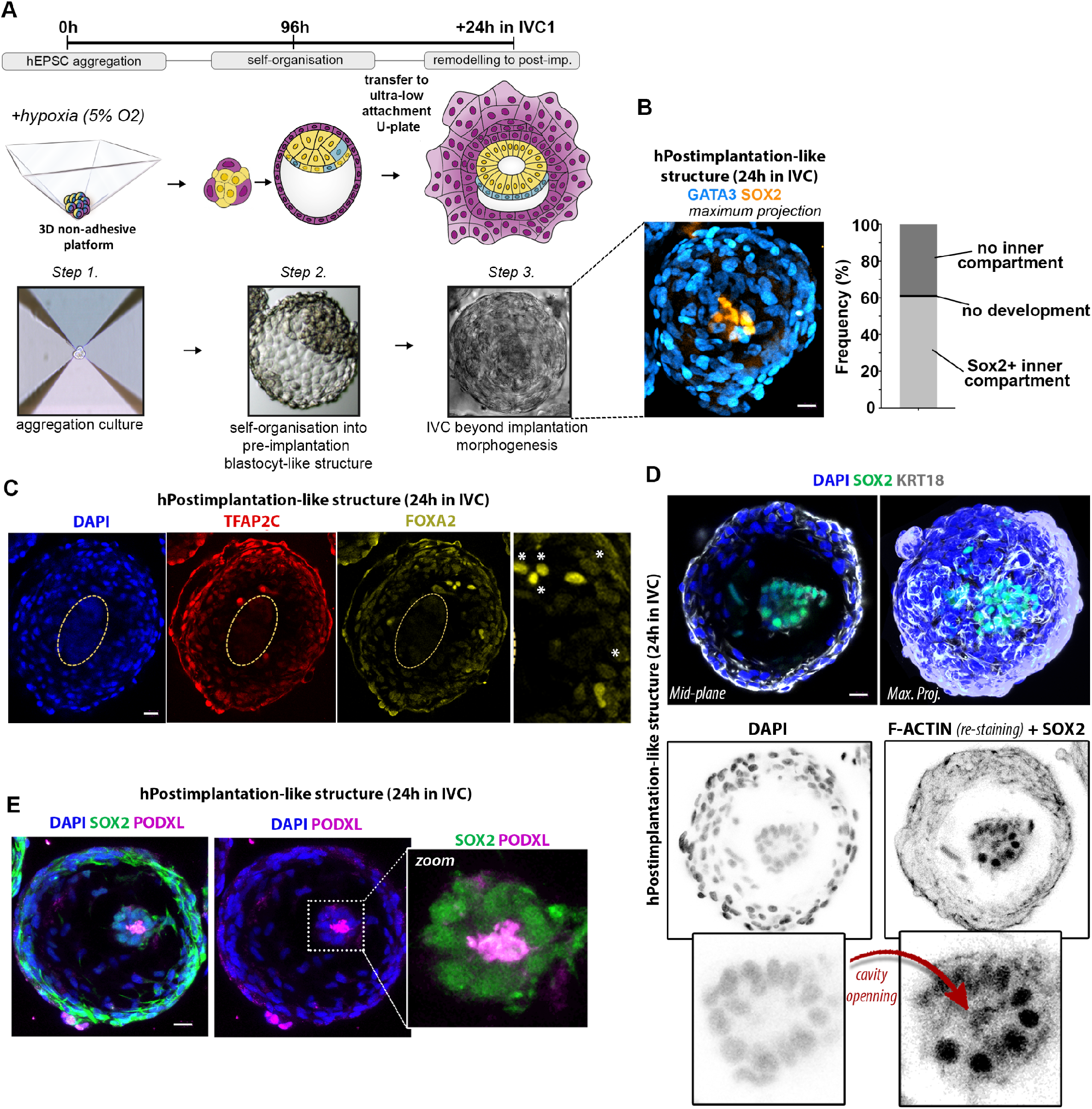
hEPSC-derived cystic structures cultured in IVC demonstrate post-implantation-like remodelling. **A.** Illustration detailing the process of *in vitro* structure formation in three steps: hEPSC aggregation, self-organisation into blastocyst-like cystic structures, and post-implantation remodelling in IVC media (see Methods). Below are phase-contrast images showing a representative structure at each of these steps. **B. Left:** Maximum projection of a representative post-implantation-like structure immunostained for GATA3 (cyan) to reveal extra-embryonic-like and SOX2 (yellow) for embryonic-like compartments. **Right:** quantification shows frequency of structures cultured in IVC media for 24h that showed SOX2+ inner compartment (60.4%, light grey), no inner compartment (38.4%, dark grey), or no development (1.2%, black). n = 260 structures scored in 3 experiments. **C.** Immunostaining showing the expression of TFAP2C (red), an extra-embryonic TE-derivative lineage marker, and FOXA2 (yellow), a Hypo marker, in a post-implantation-like structure cultured in IVC for 24h. Zoomed image on the right shows FOXA2 expressing Hypo-like cells (arrowheads). Yellow dashed-lines demarcates Epi-like inner compartment. **D.** Immunostaining of post-implantation-like structure cultured in IVC media for 24h. The top panels show a mid-plane (left) and maximum projection (right) view of a representative structure with an Epi-like compartment marked by SOX2 expression (green), surrounded by extra-embryonic compartment marked by KRT18 (white). DAPI for nuclear staining is in blue. The bottom panels show inverted images for better clarity of DAPI signal for cell nucleus on the left and F-ACTIN+SOX2 double-staining on the right. The opening of a cavity within the Epi-like compartment (nuclear SOX2 expression) is marked by F-ACTIN. n= 20 structures, 3 experiments. **E.** Immunostaining showing the formation of a amniotic-like cavity, as marked by PODXL (magenta), within the ICM, marked by SOX2 (green), of an hEPSC-derived structure after 24h culture in IVC. n= 20 structures, 3 experiments. DAPI staining is in blue. All scale bars in the figure indicate 20 um.

The restricted ability of TE lineage specification in blastocyst-like structures generated solely from hEPSCs led us to question whether these cells possess the capacity to fully generate the TE lineage in addition to Epi and Hypo commitment. We were recently able to correct a similar shortcoming in mouse blastoids by combining EPSCs with TSCs^3^. As a recent study has showed it possible to derive human TSCs from first trimester human placental samples and blastocysts^27^, this led us to test whether such TSCs could participate alongside hEPSCs to build human blastoids **(Extended Data Fig. 4a)**. To this end, we first allowed hEPSCs to aggregate for 24h after which we added hTSCs^3^. The first 24h demonstrated the increased potential to generate both embryonic Epi-like and extraembryonic Hypo-like lineages in the EP condition compared to the conventional Rset condition **(Extended Data 4b)**. Within 4 days of co-culture, we observed the formation of cystic structures having internal acentric hEPSC compartments. We also observed robust expression of blastocyst lineage markers, including GATA3, SOX2 and SOX17 **(Extended Data Fig. 4c)**. Nevertheless, it appeared that these structures struggled to form a cohesive epithelium and displayed not one but multiple cavities **(Extended Data Fig. 4c)**. Given that these hTSCs are reported to be most similar to villous cytotrophoblasts^27^, it seemed from our findings that these hTSCs may be more conducive to post- rather than pre-implantation development. We therefore tested recent conversion protocols described to transform hPSCs into putative human trophectoderm stem cells (hTESCs)^39^ **(Extended Data Fig. 4d)** that have been suggested to more closely resemble the pre-implantation TE lineage. However, we observed that these hTESCs showed little to no TE-specific marker expression, signifying an inability to maintain lineage identity upon co-culture **(Extended Data Fig. 4e)**. Thus, future work will be required to generate human TSC lines having the ability to recapitulate pre-implantation development in order to enhance this system further.

In summary, the hEPSCs we describe here show multi-directional, enhanced developmental potency enabling them to develop the morphology of human blastocysts and peri-implantation stages and, exhibit many of the molecular features of cleavage stage pre-implantation embryos. However, these cells are not the equivalent of totipotent blastomeres as although they appear able to specify Epi and Hypo cells, TE is only partially specified. This may reflect distinct molecular trajectories and an intermediate state adopted by these cells that lead to the generation of improperly differentiated cells observed in this study. Nevertheless, these cells successfully generate multicellular structures that are patterned similarly to the early human embryos. Thus, they offer the potential to be harnessed into a fully functional embryo-like platform *in vitro.* Collectively, the hEPSCs-based system we describe here recapitulates key features of natural human development from embryonic day 3 to day 9/10. It thus represents a significant step towards developing an *in vitro* model that captures the dynamics of spatiotemporal pre/peri-implantation development in a 3D context. We anticipate that this system will lead to a variety of future applications and will be pivotal in unravelling many of the enigmas of human developmental regulation.

## Acknowledgments

We are grateful to David Glover and Andy Cox for their helpful discussion and comments on the manuscript. We are grateful to Rachel S. Mandelbaum, Richard J. Paulson, Ali Ahmady and USC Fertility for their help and support to obtain images shown in Figure 1E; Angel Martin, Maria J. de los Santos and IVIRMA Valencia for images shown in Figure 2E. Human TSCs were kindly provided by Hiroaki Okae and Takahiro Arima (Tohoku University Graduate School of Medicine, Japan). The Zernicka-Goetz Lab is funded by the Welcome Trust (098287/Z/12/Z), the NIH Pioneer Award Fund (DP1 HD104575-01), Open Philanthropy/Silicon Valley Community Foundation, Weston Havens Foundation and Shurl and Kay Curci Foundation to M.ZG. B.S. is now funded by Yale School of Medicine start-up award.

## Author Contributions

B.S. designed and performed experiments and analysed the data with the help of V.J. M.Z. provided data on human embryos. T.C performed conversion protocol for hPSCs to hTESCs. B.S., V.J. and M.ZG wrote the manuscript. B.S and M.ZG. conceived and supervised the project.

---

**Extended Data Figure 1.**
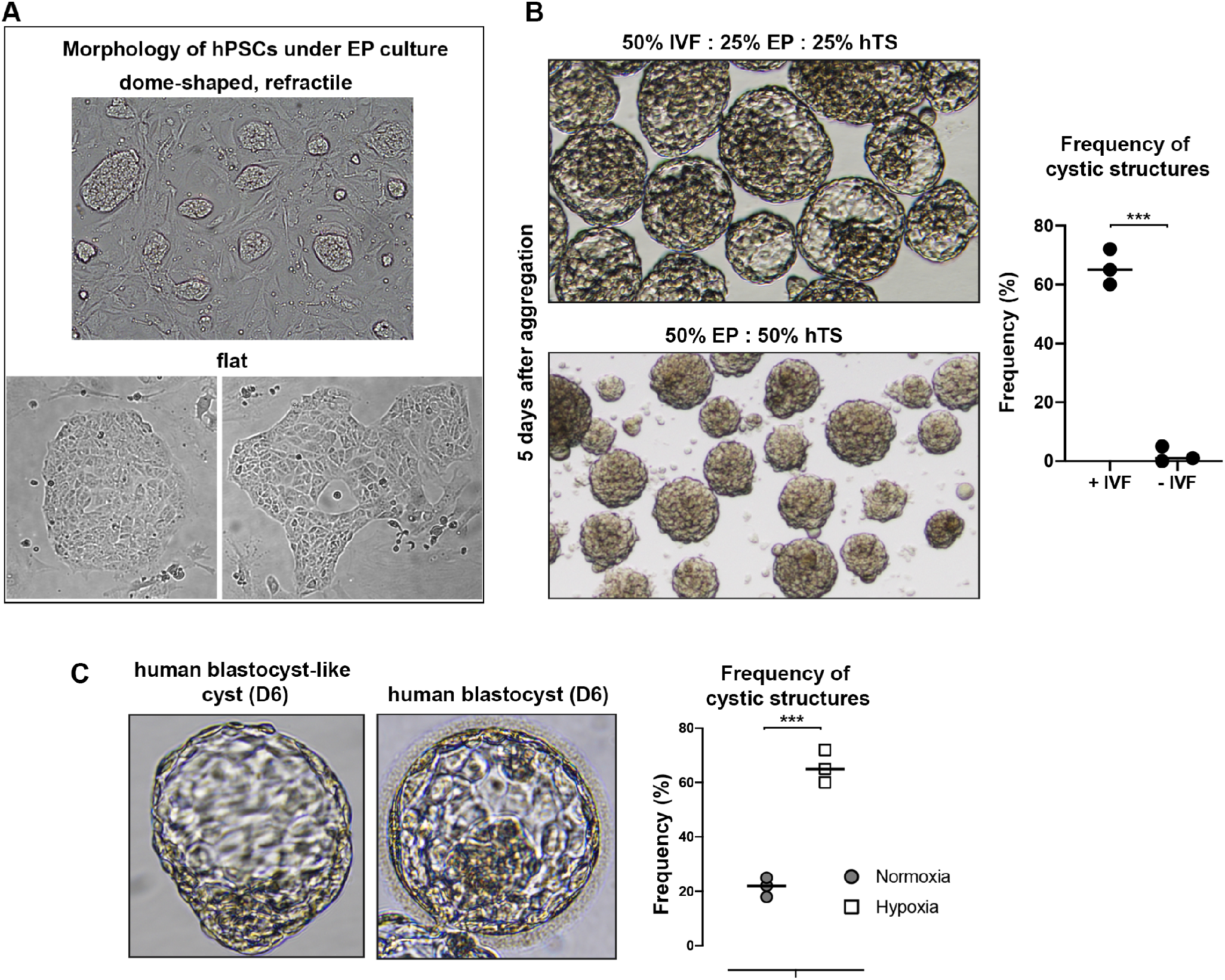
Characterisation of cellular and structural morphologies under 2D and 3D culture conditions. **A.** DIC images showing varied colony morphology of hPSCs under EP culture conditions. The top image shows dome-shaped morphology, characteristic of pluripotent cells at the naïve state, while the bottom images show flat colonies, characteristic of cells in the primed state. **B.** DIC images of hEPSC-derived cystic structures grown for 5 days with (top) and without (bottom) IVF media. Quantification on the right shows the frequency of cystic structure formation in each condition (+/-IVF). Student’s t-test; p<0.001; 3 experiments. **C.** Comparison of hEPSC-derived blastocyst-like structures at day 6 (D6) of 3D culture to the natural human blastocyst at the same developmental time point. Quantification on the right shows the frequency of cystic structure formation under 20% O_2_ (Normoxia; grey circle) and 5% O_2_ (Hypoxia; white square) conditions. Student’s t-test; p<0.001; 3 experiments. All scale bars in the figure indicate 20 um.

**Extended Data Figure 2.**
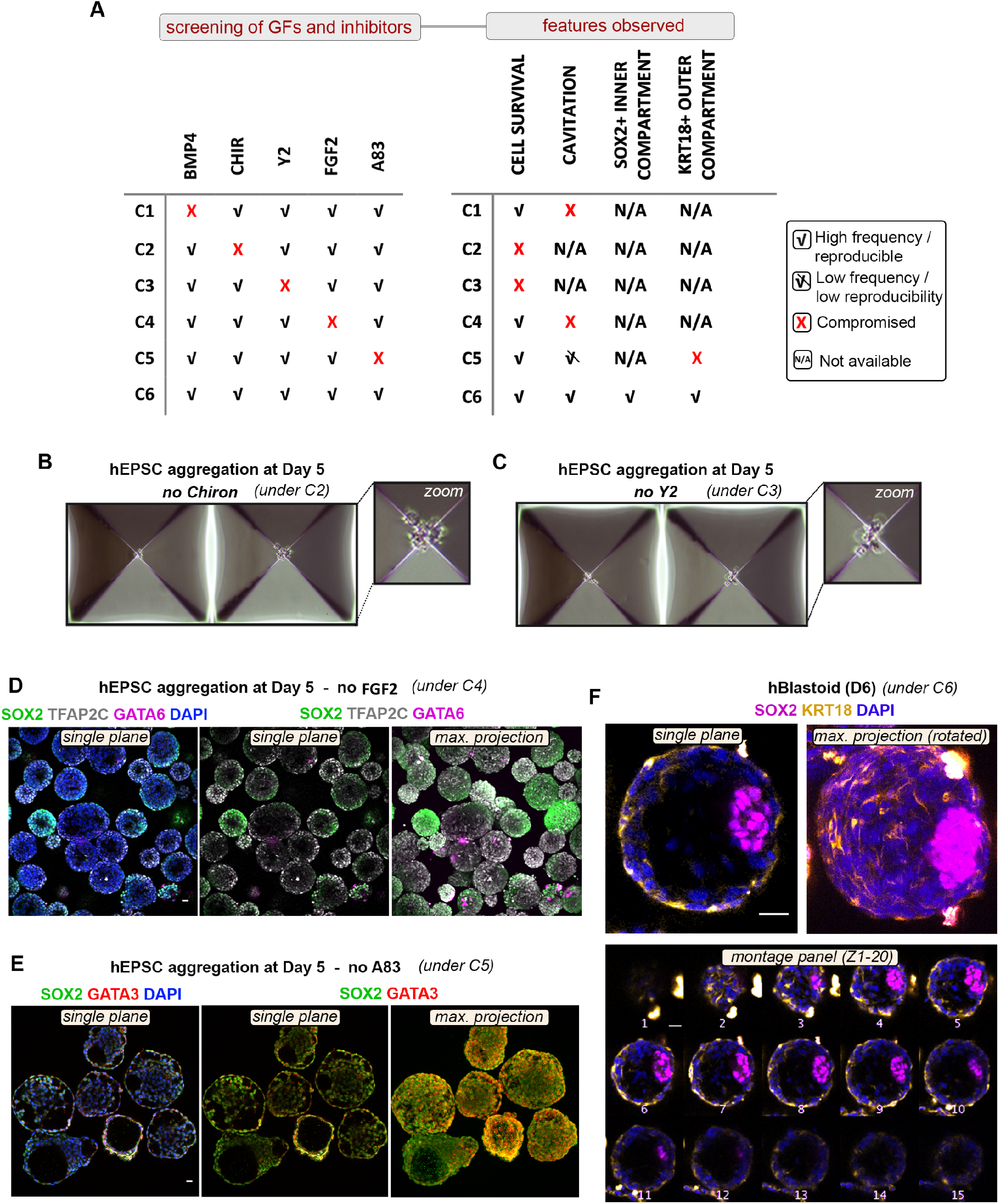
Screening of growth factors and inhibitors on hEPSC-derived structures. **A.** Table outlining tested media conditions with and without specific growth factors and inhibitors (left), and observed features from each of these conditions (right). **B, C.** Images show failure of cell survival grown for 5 days without CHIR99021 (Chiron, condition 2, C2) or Y-27632 (Y2; condition 3, C3), respectively. **D.** Immunostaining showing expression of TFAP2C (white), and GATA6 (magenta) in structures grown under condition 4 (C4) for 5 days. DAPI shows nuclear staining in blue. **E.** Immunostaining showing expression of GATA3 (red), and SOX2 (green) in structures grown under condition 5 (C5) for 5 days. DAPI shows nuclear staining in blue. **F.** Immunostaining for SOX2 (magenta), and KRT18 (yellow) in representative structure grown under condition 6 (C6) until Day 6. Images are shown in a single plane view (top left), maximum projection (top right), and as a montage panel of Z-stack slice 1-20. All scale bars in the figure indicate 20 um.

**Extended Data Figure 3.**
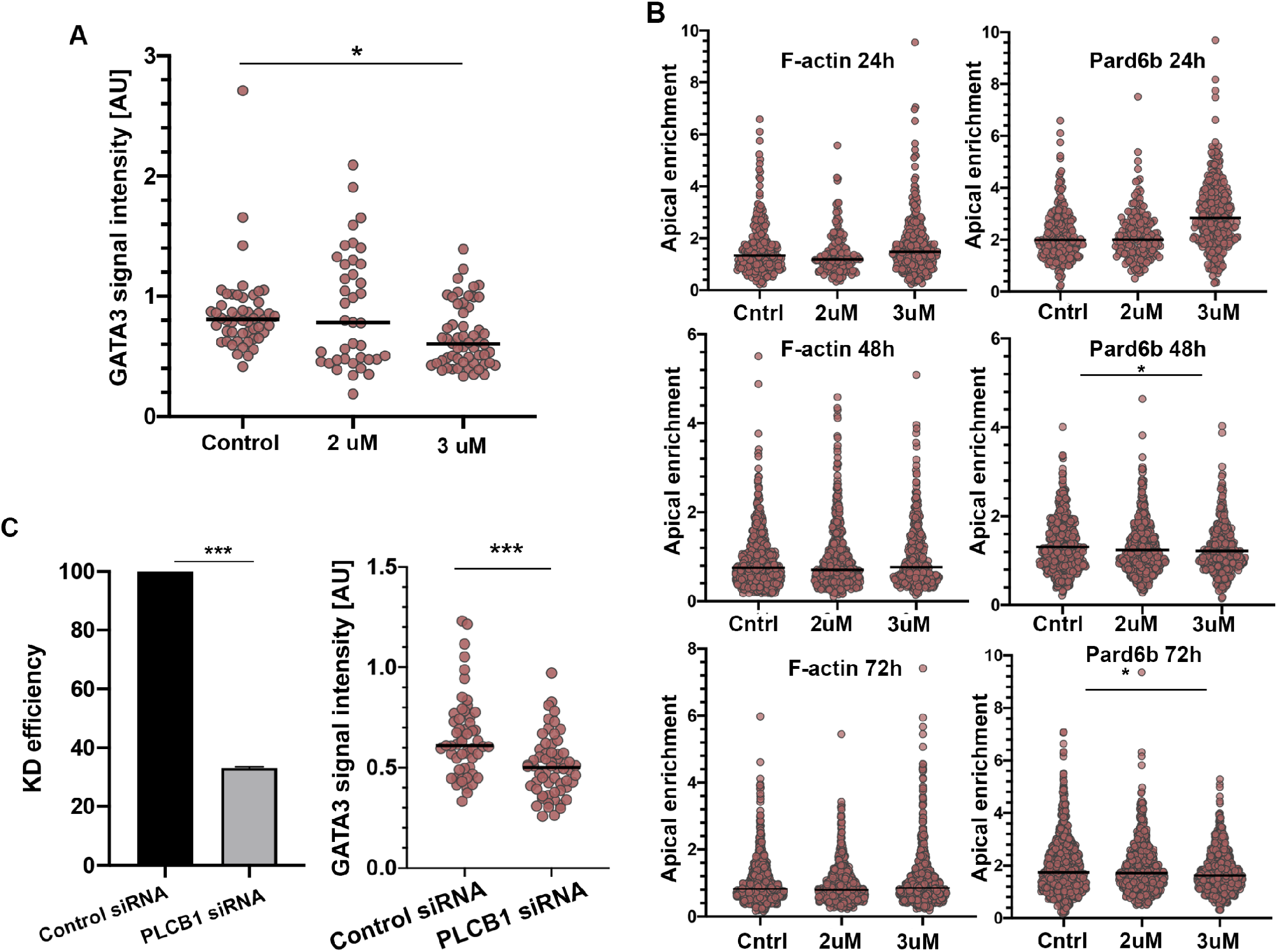
Expression of nuclear GATA3 intensity upon PLC-Protein Kinase C (PKC) pathway suppression. **A.** Quantification of nuclear GATA3 signal intensity in control and experimental groups treated with either 2uM or 3uM of selective PKC inhibitor, U73122. All measurements normalized to DAPI. Each dot represents one analysed cell. *p<0.05, ANOVA with a multiple comparisons test. Data is shown as mean S.E.M. 3 experiments. **B. F.** Apical enrichment quantification of F-ACTIN and PARD6b at 24, 48h and 72h in multicellular structures with or without addition of PLC inhibitor (U73122). Control groups received no inhibitor, while the two experimental groups were treated with either 2uM or 3uM U73122. Each dot represents one analysed cell. *p<0.05, Kruskal-Wallis test with a multiple comparisons test. Data is shown as mean S.E.M. n= 3 experiments. **C. Left:** Quantification of PLCB1 knock-down efficiency in groups treated with either control siRNA or *PLCB1* siRNA as determined by RT-PCR. Values were normalized against GAPDH. *p<0.001, Student’s t-test. n = 3 replicates. **Right:** Quantification of nuclear GATA3 signal intensity in groups treated with either Control siRNA or *PLCB1* siRNA. All measurements normalized to DAPI. Each dot represents one analysed cell. *p<0.001, Student’s t-test.

**Extended Data Figure 4.**
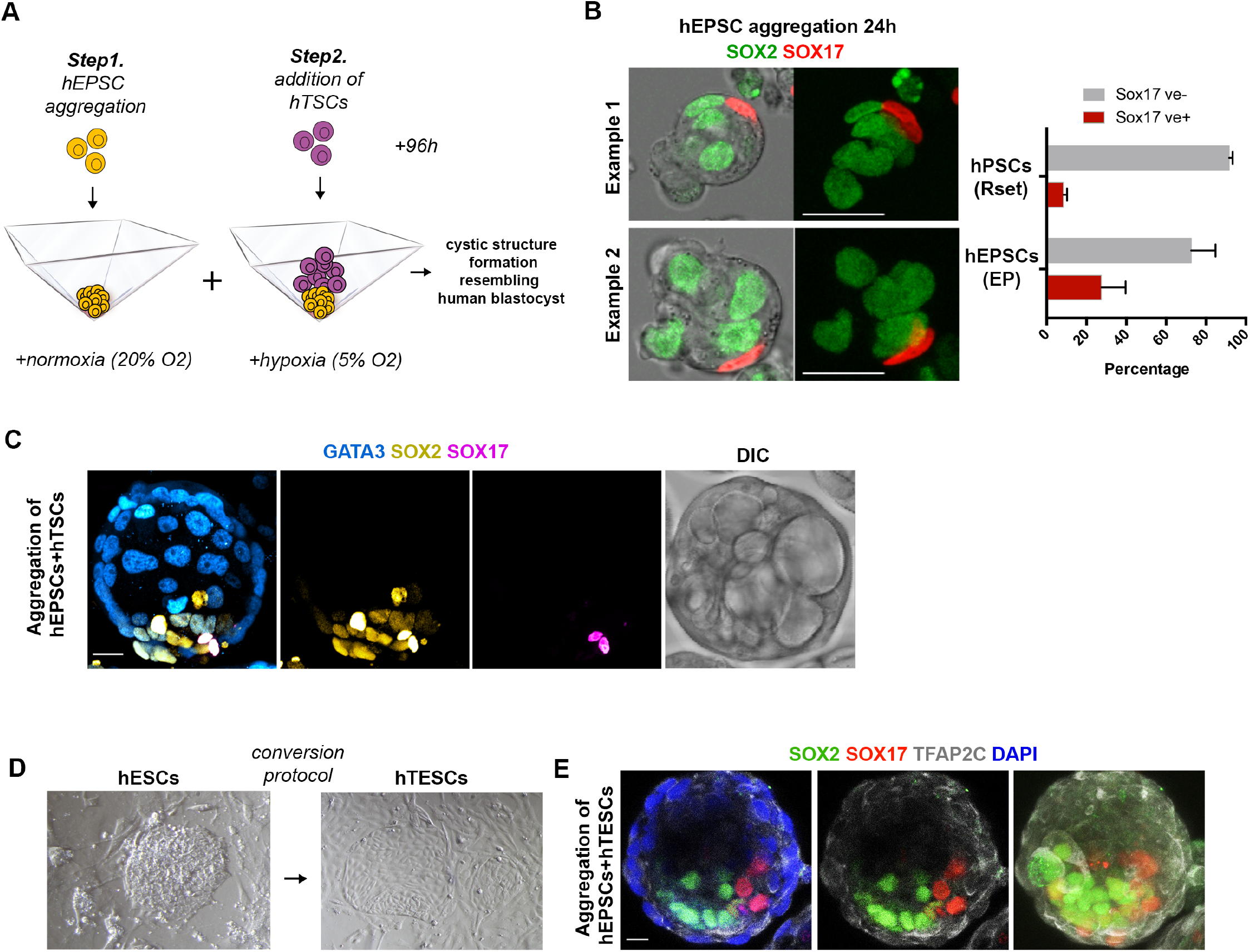
Generation of blastocyst-like structures using hEPSCs and hTSCs. **A.** 2-step protocol for generating blastocyst-like structures from EPSCs combined with hTSCs. **B.** Aggregates co-expressing endogenous SOX2 (green) and SOX17 (red) after the first step (24h) of aggregation. SOX2 indicates naive pluripotent Epi-like cells; SOX17 indicates extra-embryonic hypoblast-like cell formation. Quantification on the right shows structures scored as positive for hypoblast-like cell in conventional naïve Rset condition and EP condition. n = 100 aggregates per group. **C.** A representative structure generated from EPSCs combined with hTSCs stained for GATA3, SOX2 and SOX17. DIC image reveals the failure for cavitation. **D.** DIC image for a representative dome-shaped naïve human ESC colony in 2D culture (left) and the morphology after their conversion to human TESCs (hTESCs). **E.** Representative structure that generated from hEPSCs combined with hTESCs reveals the failure for TE-identity. SOX2 (green) for Epi-like, SOX17 (red) for Hypo-like, TFAP2C (grey) for TE-like lineage formation.

## Methods

### Data reporting

No statistical methods were used to predetermine sample size. The experiments were not randomized and the investigators were not blinded to allocation during experiments and outcome assessment.

### Ethics statement

Stem cell-derived multicellular structures described in this study show no evidence of germ line patterning, thus they do not have human organismal form or potential. Additionally, all experiments were terminated by no later than day 8 in vitro. Our research was subject to review and approval from the Human Embryo and Stem Cell (HESC) Committee of California Institute of Technology, in compliance with the ISSCR 2016 guidelines.

### Human Cell lines

The hPSC lines utilized in this study include: RUES2-RLG (kindly provided by Ali Brivanlou, The Rockefeller University, US), H9 (kindly provided by Juan Carlos Izpisua Belmonte, The Salk Institute, US) and ESI0017 (kindly provided by Juan Michael Elowitz, California Institute of Technology, US). Human TSCs were kindly provided by Hiroaki Okae and Takahiro Arima (Tohoku University Graduate School of Medicine, Japan). Each of these cell lines was tested negative for mycoplasma contamination, which was monitored on a bi-monthly basis (MycoScope™ PCR Mycoplasma Detection Kit, Genlantis).

### Cell culture

All hEPSC lines were maintained under 20% O2 and 5% CO2 at 37°C conditions on irradiated CF1 mouse embryonic fibroblasts (MEF) feeder cells. hEPSCs were grown using ‘human Expanded Potential’ (hEP) medium consisting of DMEM/F12 (Thermo Fisher Scientific, 11320-033), Neurobasal-A (Thermo Fisher Scientific, 21103-049), N2 supplement (Thermo Fisher Scientific, 17502-048), B27 supplement (Thermo Fisher Scientific, 12587-010), 1% GlutaMAX (Thermo Fisher Scientific, 35050-061), 1% nonessential amino acids (Thermo Fisher Scientific, 11140-050), 0.1 mM b-mercaptoethanol (Thermo Fisher Scientific, 31350-010), penicillin-streptomycin (Thermo FisherScientific, 15140–122) and 5% knockout serum replacement (KSR, Thermo Fisher Scientific, A3181502). LCDMYI supplementation was added as indicated at the following concentrations: 10 ng ml−1 recombinant human LIF (L, 10 ng ml−1; Peprotech, 300-05), CHIR99021 (C, 1 mM; Stem Cell Technologies), (S)-(+)-Dimethindenemaleate (D, 1 mM; Tocris, 1425) and Minocycline hydrochloride (M, 2 mM; Santa Cruz Biotechnology, sc-203339), 1. All hEPSCs were used before reaching P70 and cell cultures were examined by eye to monitor for spontaneous differentiation of colonies into mesenchymal-like cells.

hTSCs were cultured on 6-well plates pre-coated with 5 mg/ml Col IV at 37C for at least one hour, as previously described in Okae *et al^27^.* Cells were grown in ‘human Trophoblast stem cell’ (hTS) medium consisting of DMEM/F12 supplemented with 0.1 mM b-mercaptoethanol, 0.2% FBS, 0.5% Penicillin-Streptomycin, 0.3% BSA (A8806-5G, Sigma-Aldrich), 1% ITS-X supplement (51500-056, Thermo Fisher Scientific), 1.5 mg/ml L-ascorbic acid (A4403, Sigma-Aldrich), 50 ng/ml EGF (62253-63-8, Sigma-Aldrich), 2 mM CHIR99021, 0.5 mM, A83-01 (72024, Stemcell Technologies), 1 mM SB431542 (Stem Cell Technologies), 0.8 mM VPA (Sigma-Aldrich) and 5 mM Y27632 hTESCs were grown on 6-well plates as described in Mischeler *et al.* ^39^ with some modifications. Plates were pre-coated with coated with 3 μg/ml of vitronectin (07180, Stemcell Technologies) and 5mg/mL Col IV (C7521-5MG, Sigma-Aldrich), instead of laminin 521 and cells were grown in TM4 media consisteing of the following: TeSR-E6 medium supplemented with CYM5541 (2 μM), A 83-01 (0.5 μM), FGF10 (25ng/ml) and CHIR99021 (2 μM).

### Preparing and Plating Cell Suspensions for ‘‘AggreWell’’ Aggregation Experiments

AggreWell 400 format plates were prepared following the manufacturer’s protocol. Briefly, wells were rinsed with the rinsing solution (Stem Cell Technologies), centrifuged for 5 min at 2000g and incubated at room temperature in the tissue culture hood for 20 min. The wells were then washed with 2 ml of 1X PBS. After PBS removal, 500 ml of final culture medium (*IVF-hEP-hTS*, see below) was added to each well and the plate placed at 37C and 5% CO2 until ready to use.

### Generation of multicellular aggregates in 3D

To begin, hEPSCs were dissociated to single cells by incubation with Accutase (07920, Stem Cell Technologies) at 37°C for 3 min. Cells were collected and pelleted by centrifugation for 4 min at 1000 rpm and resuspended in hEP-LCDMYI medium (described above). This cell suspension was pre-incubated at 37°C in an atmosphere of 5% CO_2_ on gelatinized tissue-culture-grade plates for 30 min to remove inactive MEFs.

Post incubation on gelatin plates, cells were counted using a haemocytometer and a total of 7200 hEPSCs was added to 1mL of media composed of 50% *IVF* media (Continuous Single Culture-NX Complete (CSCM-NXC)) (90168, FUJIFILM), 25% *hEP* media, and 25% *hTS* media. This media was also supplemented with CHIR99021 (2uM), Y27632 (5uM), BMP4 (20ng/mL), FGF2 (40ng/mL), and A83-01 (2uM). Cell suspensions were added dropwise to the Aggrewells. All wells without cells were filled with 1mL PBS to humidify the local atmosphere to minimize evaporation. The AggreWell plate was then centrifuged for 3 min at 100g, and placed at 37°C under hypoxic conditions (5% CO_2_ and 5% O_2_). After 48h, media was removed from wells and replaced with fresh culture media as described above, although FGF2 concentration was lowered to 20ng/mL and A83-01 was omitted. Cells were left to grow for an additional 48-72h until proper morphology was observed, at which point structures were fixed for immunostaining or transferred to IVC media (see below) for mimicking development beyond implantation stages.

### Criteria for selecting multicellular aggregates structures

Following completion of any given aggregation experiment (from day 4 to 6), all cystic structures those clearly displaying a cavity were included in further analyses. Non-cavitated structures were excluded from downstream analyses.

### Co-culture of hEPSCs with hTSCs/hTESCs

To perform two-step aggregation experiments, hEPSCs were first seeded as described above hEP-LCDMYI media. After a 24h period of aggregation, hTSC or hTESC colonies were dissociated to single cells, and counted using a haemocytometer. For hTSC experiments, 50% *hEP* media and 50% *hTS* media is used as described above. For hTESC experiments, hTS media was replaced with TeSR-E6, all other factors remained the same. A total of 16,800 hTSCs/hTESCs were added per well and the plate was placed at 37°C, 5% CO2 and 5% O2.

### *In Vitro* Culture of hEPSC blastocyst-like structures beyond implantation

To prepare plate for *in vitro* culture, 150uL of modified *IVC1* (*mIVC1*) media was added to each well of a 96-well ultra-low attachment U-shaped plate (7007, Costar). *mIVC1* media consisted of the following: Advanced DMEM/F12 (12634-010; Thermo Fischer Scientific; Waltham, US) supplemented with 20% (vol/vol) heat-inactivated FBS (16141079, Thermo Fisher Scientific), 2mM GlutaMAX, penicillin (25 units/ml)/Streptomycin (25 μg/ml), 1X ITS-X (10 mg/L insulin, 5.5 mg/L transferrin, 0.0067 mg/L sodium selenite, 2 mg/L etholamine; 51500-056; Thermo Fisher Scientific; Waltham, US), 8 nM β-estradiol (E8875; Sigma-Aldrich; St. Louis, US), 200 ng/ml progesterone (P0130; Sigma-Aldrich; St. Louis, US), 25 μM *N*-acetyl-_L_-cysteine (A7250; Sigma-Aldrich; St. Louis, US), 17nm IGF1, 20ng/mL FGF2 (Gibco), FGF4 (25 ng/mL; R&D Systems, 5846-F4) and heparin (1 mg ml^−1^; Sigma, H3149).

### Immunofluorescence Staining

Stem cell-derived structures were fixed in 4% paraformaldehyde (Electron Microscopy Sciences, 15710) for 20 min at room temperature, and then washed twice in PBT [phosphate-buffered saline (PBS) plus 0.05% Tween-20]. Structures were permeabilized for 30 min at room temperature in PBS containing 0.3% Triton-X-100 and 0.1% glycine. Primary antibody incubation was performed overnight at 4°C in blocking buffer [PBS containing 10% fetal bovine serum (FBS), 1% Tween-20]. The following day, embryos were washed twice in PBT, then incubated overnight at 4°C with secondary antibody (1:500) in blocking buffer. Structures were washed twice in PBT buffer and then transferred to PBT drops in oil-filled optical plates before confocal imaging. The antibodies used are given in **Supplementary Data Table 1**.

For human embryo images shown in Figure 2e, embryos were fixed in IVIRMA Valencia, washed twice in a PBS solution containing 0.1% Tween-20 (Sigma, cat. no. P9416) and immediately placed into a 0.5 ml PCR tube within an oil-PBS-oil interphase. Tubes were stored at 4°C were shipped to the University of Cambridge for immunofluorescence.

### Image Data Acquisition, Processing, and Quantification

Fluorescence images were acquired with an inverted Leica SP8 confocal microscope (Leica Microsystems), using a Leica Fluotar VISIR 0.95 NA 25x objective. Fluorophores were excited with a 405-nm diode laser (DAPI), a 488-nm argon laser (GFP), a 543-nm HeNe laser (Alexa Fluor-543/555) and a 633-nm HeNe laser (Alexa Fluor-633/647). Images were acquired with 0.5–1.2 mm z-separation. Raw data were processed using open-source image analysis software ‘‘Fiji’’ software and assembled in Photoshop CC 2019 (Adobe). Digital quantifications and immunofluorescence signal intensity graphs were obtained using Fiji software.

*Apical enrichment analysis:* F-actin and PARD6 polarisation were measured in a single focal plane, by taking the middle plane of the aggregate. A freehand line of the width of 0.5μm was drawn along the cell-contact free surface (apical domain), or cell-contact (basal) area of the cell, signal intensity was obtained via the Region of Interest (ROI) function of Fiji. The apical/basal signal intensity ratio is calculated as: I(apical)/I(basal). A cell is defined as polarised when the ratio between the apical membrane and the cytoplasm signal intensity exceeds 1.5.

*GATA3 expression analysis:* the nucleus of each cells is masked using the Region of Interest (ROI) tool of Fiji. The average signal intensity of the ROI is calculated and a cell is defined as GATA3 positive when the nucleus to cytoplasm signal intensity exceeds 1.5.

### siRNA-mediated knock-down in hEPSC-derived aggregates

Transfections of siRNA were performed using Lipofectamine RNAi MAX (13778075, Thermo Fisher Scientific) according to the manufacturer’s instructions. Upon seeding hEPSCs into AggreWells (as described above), Lipofectamine and siRNA (Qiagen, Hs_PLCB1_4, SI00115521; Qiagen, Hs_PLCB1_6, SI02781184; Qiagen, negative control siRNA, 1022076) against target genes with Opti-MEM (31985070, Gibco) is mixed and the mixture of either control siRNA or PLCB1 siRNA were evenly added into each well. Cell aggregates at 48h were collected to analyse the gene expression by qRT-PCR.

### Bulk qRT-PCR Analysis

Total RNA was extracted with using Arcturus PicoPure™ RNA Isolation Kit (12204-01, Applied Biosystems) as per manufacturer’s instructions. QRT-PCR was performed with the Power SYBR Green RNA-to-CT 1-Step Kit (Life Technologies) and a Step One Plus Real-time PCR machine (Applied Biosystems). The amounts of mRNA were measured with SYBR Green PCR Master Mix (Ambion). Relative levels of transcript expression were assessed by the ΔΔCt method, with Gapdh as an endogenous control. For qPCR primers used, see **Supplementary Data Table 2**.

### Statistics and Reproducibility

Statistical tests were performed on GraphPad Prism 8.0 software. Where appropriate, Student’s t-tests (two groups) or analysis of variance (multiple groups) were performed. Figure legends indicate the number of independent experiments performed in each analysis. Unless otherwise noted, each experiment was performed at least two times.

---

**Extended Data Table 1.** List of primary antibodies.

| Targeted Protein | Species | Dilution | Catalog No. | Vendor |
| --- | --- | --- | --- | --- |
| **PARD6** | Rabbit | 1:200 | sc-166405 | Santa Cruz Biotechnology |
| **E-Cadherin (Clone 36)** | Mouse | 1:200 | 610182 | BD Biosciences |
| **OCT3/4** | Rabbit | 1:200 | Sc-9081 | Santa Cruz Biotechnology |
| **SOX2** | Rabbit | 1:200 | D6D9 | Cell Signaling Technologies |
| **GATA3** | Goat | 1:200 | AF2605 | R&D Systems |
| **PODXL** | Mouse | 1:20 | MAB1658 | R&D Systems |
| **SOX17** | Goat | 1:200 | AF1924 | R&D Systems |
| **FOXA2** | Rabbit | 1:200 | D56D6 | Cell Signaling Technologies |
| **GATA6** | Goat | 1:200 | AF1700 | R&D Systems |
| **Alexa Fluor® 488 Phalloidin (F-ACTIN)** | N/A | 1:500 | A12379 | Thermo Fisher Scientific |
| **TFAP2C** | Goat | 1:200 | AF5059 | R&D Systems |
| **KRT18** | Mouse | 1:200 | Ab668 | Abcam |
| **Anti-GFP** | Rat | 1:1000 | 04404-84 | Nacalai Tesque |

**Extended Data Table 2.**
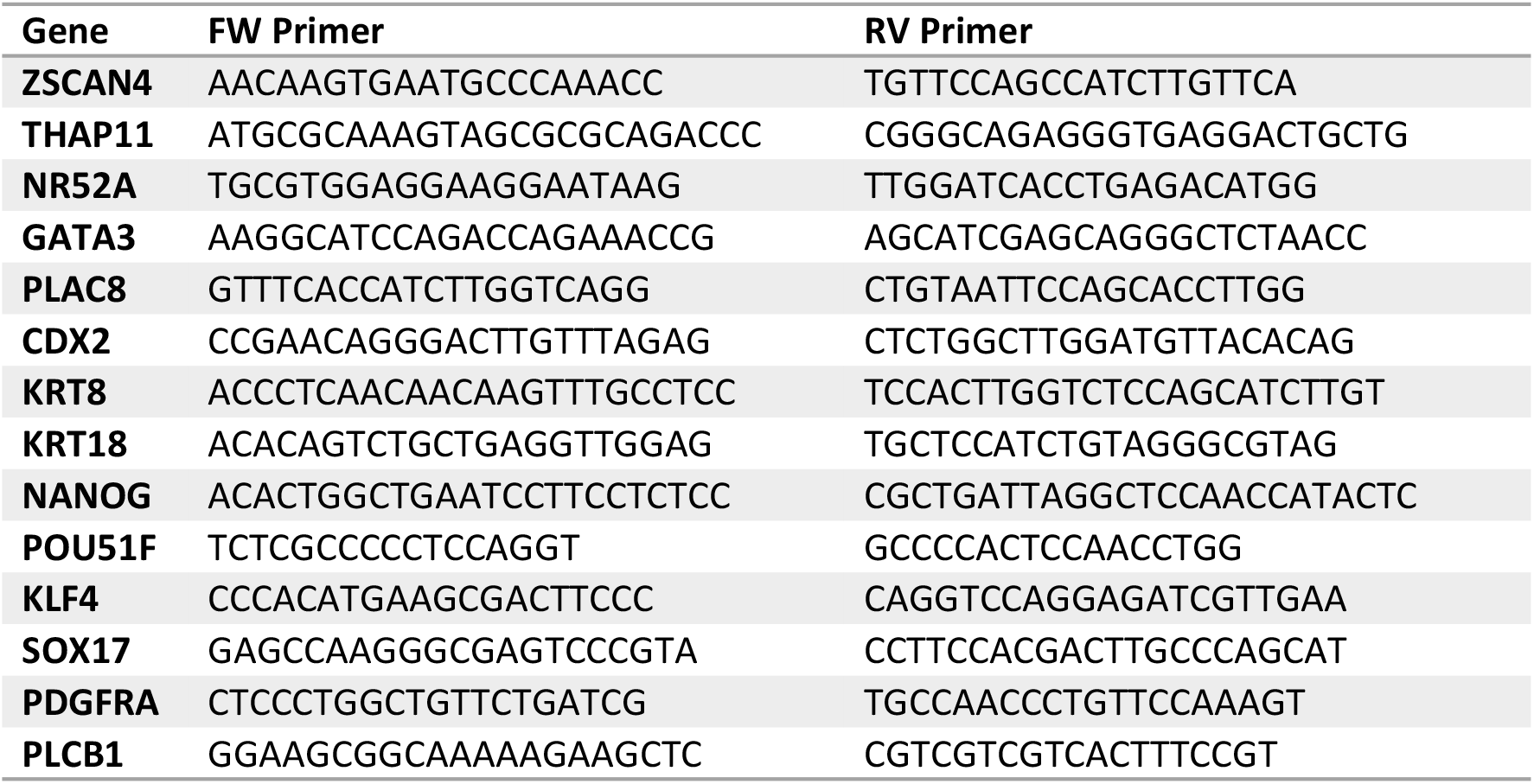
List of primers for RT-qPCR.

